# Toward leveraging big data in human functional connectomics: Generalization of brain graphs across scanners, sessions, and paradigms

**DOI:** 10.1101/160440

**Authors:** Hengyi Cao, Sarah C. McEwen, Carrie E. Bearden, Jean Addington, Bradley Goodyear, Kristin S. Cadenhead, Heline Mirzakhanian, Barbara A. Cornblatt, Doreen M. Olvet, Daniel H. Mathalon, Thomas H. McGlashan, Diana O. Perkins, Aysenil Belger, Larry J. Seidman, Heidi Thermenos, Ming T. Tsuang, Theo G.M. van Erp, Elaine F. Walker, Stephan Hamann, Scott W. Woods, Tyrone D. Cannon

## Abstract

While graph theoretical modeling has dramatically advanced our understanding of complex brain systems, the feasibility of aggregating brain graphic data in large imaging consortia remains unclear. Here, using a battery of cognitive, emotional and resting fMRI paradigms, we investigated the reproducibility of functional connectomic measures across multiple sites and sessions. Our results revealed overall fair to excellent reliability for a majority of measures during both rest and tasks, in particular for those quantifying connectivity strength, network segregation and network integration. Higher reliabilities were detected for cognitive tasks (vs rest) and for weighted networks (vs binary networks). While network diagnostics for several primary functional systems were consistently reliable independently of paradigm, those for cognitive-emotional systems were reliable predominantly when challenged by task. Different data aggregation approaches yielded significantly different reliability. In addition, we showed that after accounting for observed reliability, satisfactory statistical power can be achieved in the multisite context with a total sample size of approximately 250 when the effect size is at least moderate. Our findings provide direct evidence for the generalizability of brain graphs for both resting and task paradigms in large consortia and encourage the use of multisite, multisession scans to enhance power for human functional connectomic studies.

## Introduction

Since its debut in the last decade (Sporns O et al. 2005), the study of the human functional connectome has become an increasingly appealing research frontier in modern neuroscience. The brain connectome is typically modelled using graph theoretical methods, which decompose the functional architecture of the brain into a large set of nodes and interconnecting edges (Bullmore ET and DS Bassett 2011). This approach has greatly advanced our understanding of the functional organization of the brain, bringing valuable insights into the topological characteristics of brain systems (Power JD et al. 2011) and variations therein related to neural development (Fair DA et al. 2009), aging (Meunier D et al. 2009), and clinical brain disorders (Buckner RL et al. 2009; Lynall ME et al. 2010; Cao H *et al*. 2016).

We are now in the era of “big data”, where large research consortia have been established around the world and hundreds or thousands of imaging scans could potentially be pooled to pursue questions that can only be addressed with large sample sizes (Biswal BB et al. 2010). Such applications include ascertaining genetic determinants of brain network structure (Richiardi J et al. 2015) or elucidating patterns in brain network architecture predictive of low-incidence disease among individuals at risk (Cao H *et al*. 2016). However, while moderate to high test-retest reliability of brain graph properties have been demonstrated in both resting state (Braun U *et al*. 2012; Cao H *et al*. 2014; Termenon M *et al*. 2016) and cognitive tasks (Cao H *et al*. 2014) using data acquired on a single scanner, it remains unclear whether the increased sample size associated with pooling data collected across different scanners is offset by attenuated reliability of network analysis measures in relation to statistical power. The utility of big data fusion in human functional connectomics will be constrained by the answer to this question.

Here, using the data from the North American Prodrome Longitudinal Study (NAPLS) consortium (Addington J et al. 2012), we examined the feasibility of aggregating multisite, multisession functional magnetic resonance imaging (fMRI) data in the study of brain graphs. In this work, eight subjects were scanned twice (on consecutive days) at each of the eight study sites across the United States and Canada using a battery of five fMRI paradigms including four cognitive tasks and a resting state scan. This unique sample allows us to explicitly answer the question of whether it is feasible (i.e., achieving acceptable levels of reliability) to aggregate fMRI data acquired from multiple sites and sessions and to determine which approach to aggregating such data maximizes reliability. Generalizability theory was used to quantify reliability of graph theoretical metrics, first for the full eight-site, two-session study, and then for the circumstance in which a given subject is scanned once on one scanner drawn randomly from the set of all available scanners (i.e., paralleling the design of the typical “big data” study in which scans from a single session are pooled across multiple sites). We compared reliability of graph theoretical metrics across the five fMRI paradigms and across four different graph construction schemes and isolated the most reliable nodes in the brain for each paradigm. We also estimated the required sample size to achieve satisfactory statistical power in a multisite study and investigated the effects of two data pooling methods (“merging raw data” and “merging results”) on the reliability of the resulting brain graphs. The results of this study provide evidence for the feasibility, sample size determination and optimal method of pooling large sets of graph theoretical measures for functional connectomics research in large consortia.

## Methods

### Subjects

A sample of eight healthy traveling subjects (age 26.9 ± 4.3 years, 4 males) was included as part of the North American Prodrome Longitudinal Study (NAPLS-2) consortium (Addington J et al. 2012). The consortium comprises eight study sites across the United States and Canada: Emory University, Harvard University, University of Calgary, University of California Los Angeles (UCLA), University of California San Diego (UCSD), University of North Carolina Chapel Hill (UNC), Yale University, and Zucker Hillside Hospital (ZHH). Each site recruited one subject and the participants traveled to each of the eight sites in a counterbalanced order. At each site, subjects were scanned twice on two consecutive days with the same fMRI paradigms, resulting in a total of 128 scans (8 subjects × 8 sites × 2 days) for each paradigm. All scans were completed within a period of two months, during which time no changes were made to the MRI scanners at each site.

All participants received the Structured Clinical Interview for Diagnostic and Statistical Manual of Mental Disorders (DSM-IV-TR (First MB et al. 2002)) and Structured Interview for Prodromal Syndromes (McGlashan TH et al. 2001), and were excluded if they met the criteria for psychiatric disorders or prodromal syndromes. Other exclusion criteria included a prior history of neurological or psychiatric disorders, substance dependency in the last six months, IQ < 70 (assessed by the Wechsler Abbreviated Scale of Intelligence (Wechsler D 1999)) and the presence of a first-degree relative with mental illness. All subjects provided informed consent for the study protocols approved by the institutional review boards at each site.

### Experimental paradigms

The NAPLS-2 consortium included a battery of five paradigms targeting functional domains of interest in cognitive neuroscience: a verbal working memory paradigm (hereafter WM paradigm), a paired-associates encoding paradigm for episodic memory (hereafter EM encoding paradigm), a paired-associates retrieval paradigm for episodic memory (hereafter EM retrieval paradigm), a facial emotional processing paradigm (hereafter EP paradigm) and a resting-state paradigm (hereafter RS paradigm). These paradigms have been described in detail in previous studies (Forsyth JK et al. 2014; Gee DG et al. 2015; Noble S et al. 2016) but are summarized briefly below.

The WM paradigm is a block-designed Sternberg-style task where subjects viewed a set of uppercase consonants (each set displayed for 2 s, followed by a fixation cross for 3 s). After each set, a lowercase probe appeared and the participants were instructed to indicate if the probe matched any of the consonants from the previous set by pressing designated buttons. Four conditions were presented in the task targeting four working memory loads with 3, 5, 7 and 9 consonants in the target sets. Each load comprised a total of 12 trials with 50% matched trials. The resting-state fixation blocks were interspersed throughout the task to provide a baseline. The entire task lasted for 9 min (184 whole-brain volumes).

The EM encoding task used an event-related paradigm where subjects were presented a series of semantically unrelated word pairs for objects from 12 different categories (e.g., animals, transportations, food, etc.) and colored picture pairs depicting each word. During each trial, participants were asked to imagine the two objects interacting together and then pressed a button once a salient relationship had been built between the two words. Each trial was displayed for 4 s and followed by a jittered inter-stimulus interval between 0.5-6 s. In the active baseline condition, subjects were presented by a series of one-digit number pairs and colored squared pairs. Participants were asked to sum up the two numbers and press a button once the summation had been calculated. The paradigm consisted of 32 encoding trials and 24 baseline trials and lasted for 8.3 min (250 whole-brain volumes).

The EM retrieval paradigm followed directly after the EM encoding task. In this task, a pair of words was presented on the screen on each trial and subjects were asked to indicate whether the given word pair had been presented during the encoding paradigm by ranking their confidence level. The retrieval paradigm consisted of 64 trials where 50% had been presented during encoding task. In the active baseline condition, participants were instructed to press the button corresponding to a confidence level presented on the screen. The retrieval run lasted for 7.3 min (219 whole-brain volumes).

The EP task consisted of two consecutive identical runs on each day. Each run comprised five conditions where subjects viewed a set of emotional faces or geometric shapes. In the emotion matching condition, participants were instructed to choose which of the two faces shown on the screen presented the same emotion as a target face. In the emotion labeling condition, subjects were asked to choose which of the two labels (e.g., angry, scared, surprised, happy) depicted a target face. In the gender matching condition, subjects needed to select which of the two faces on the screen was the same gender as a target face. In the gender labeling condition, participants selected which gender label (i.e., male or female) corresponded to a target face. In the shape matching condition, participants were asked to match two corresponding geometric shapes. Each block lasted for 50 s with 10 trials. The entire task was performed in two separate runs with 5.5 min (132 whole-brain volumes) each.

RS is a 5-min eyes-open paradigm (154 whole-brain volumes) where subjects were asked to lay still in the scanner, relax, gaze at a fixation cross, and not engage in any particular mental activity. After the scan, investigators confirmed with the participants that they had not fallen asleep in the scanner.

To ensure successful manipulation of active tasks, we checked task response rates for each scan. The scans with a response rate < 50% were excluded for data analysis. This resulted in a total of 2 scans for the EM encoding paradigm. In addition, for each of the WM, EM encoding, EM retrieval and EP tasks, 1 scan was unusable due to technical artifacts, and 1 scan for the EM encoding paradigm had shortened time series. These data were also excluded from analysis.

### Data acquisition

Imaging data were acquired from eight 3T MR scanners with three different machine models. Specifically, Siemens Trio scanners were used at Emory, Harvard, UCLA, UNC and Yale, GE HDx scanners were used at UCSD and ZHH, and a GE Discovery scanner was used at Calgary. The Siemens sites employed a 12-channel head coil and the GE sites employed an 8-channel head coil. fMRI scans were performed by using gradient-recalled-echo echo-planar imaging (GRE-EPI) sequences with identical parameters at all eight sites: 1) WM paradigm: TR/TE 2500/30 ms, 77 degree flip angle, 30 4-mm slices, 1mm gap, 220 mm FOV; 2) EM encoding and retrieval paradigms: TR/TE 2000/30 ms, 77 degree flip angle, 30 4-mm slices, 1mm gap, 220 mm FOV; 3) EP paradigm: TR/TE 2500/30 ms, 77 degree flip angle, 30 4-mm slices, 1mm gap, 220 mm FOV; 4) RS paradigm: TR/TE 2000/30 ms, 77 degree flip angle, 30 4-mm slices, 1-mm gap, 220-mm FOV. In addition, we also acquired high-resolution T1-weighted images for each participant with the following sequence: 1) Siemens scanners: magnetization-prepared rapid acquisition gradient-echo (MPRAGE) sequence with 256 mm × 240 mm × 176 mm FOV, TR/TE 2300/2.91 ms, 9 degree flip angle; 2) GE scanners: spoiled gradient recalled-echo (SPGR) sequence with 260 mm FOV, TR/TE 7.0/minimum full ms, 8 degree flip angle.

### Data preprocessing

Data preprocessing followed the standard procedures implemented in the Statistical Parametric Mapping software (SPM8,http://www.fil.ion.ucl.ac.uk/spm/software/spm8/). The same preprocessing pipelines were used for each paradigm. In brief, all fMRI images were slice-time corrected to the first slices of each run, realigned for head motion, registered to the individual T1-weighted structural images, and spatially normalized to the Montreal Neurological Institute (MNI) template with a resampled voxel size of 2×2×2 mm3. Finally, the normalized images were spatially smoothed with an 8 mm full-width at half-maximum (FWHM) Gaussian kernel.

All preprocessed images were then examined for head motion. Specifically, we quantified frame-wise displacements (FD) for each subject in each run based on the previous definition (Power JD et al. 2012). The scans with an average FD 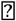 0.5 mm were shown to have a pronounced within-subject effect on connectivity (Power JD et al. 2012) and thus were discarded. This resulted in the exclusion of 1 scan for the EM encoding task, 1 scan for the EM retrieval task and 2 scans for the EP task. As a consequence, the final number of scans included for further network analysis were 127 for the WM paradigm, 123 for the EM encoding paradigm, 126 for the EM retrieval paradigm, and 125 for the EP paradigm.

### Construction of brain graphs

#### Overview of brain graph analysis

Our brain graph analysis followed closely with the standard approaches reported in the literature (Bullmore E and O Sporns 2009; Bullmore ET and DS Bassett 2011; Cao H *et al*. 2014; Cao H *et al*. 2016; Gu Q et al. 2017) and aimed to cover several different graph construction schemes. Particularly, nodes and edges are two fundamental elements in the construction of brain networks. The definitions of nodes and edges differ in the literature in terms of different brain atlases and different connection weights. While brain graphs derived from distinct processing schemes are qualitatively similar (Wang J et al. 2009; Zalesky A et al. 2010; Lord A et al. 2016) and thus might all be valid in the study of human connectomes, the comparative reliability of different schemes in the context of multisite, multisession studies is unclear. Here, we focused our analysis on two widely used brain atlases (the AAL atlas (Tzourio-Mazoyer N et al. 2002) and the Power atlas (Power JD *et al*. 2011)) and two types of graphs (binary graph and weighted graph). Consequently, four different graph models were constructed for each scan in our data: AAL binary graph, AAL weighted graph, Power binary graph and Power weighted graph. Fig. 1 provides a diagram describing the graph construction procedures.

**Fig. 1.**
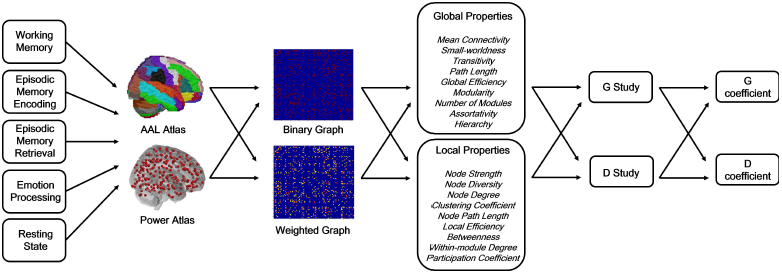
Diagram of the data processing pipeline used in the study. The time series were extracted from five fMRI paradigms using two different brain atlases, and binary and weighted brain networks were subsequently constructed from the extracted time series. Generalizability theory with both G-and D-study parameterization were employed to investigate the reliability of the graph based connectomic measures computed from these networks. See Methods for details.

#### Node definition

We used two different node definitions (anatomy-based and function-based) to construct brain graphs, in order to investigate how different atlases would influence the results. An anatomically-based definition was given by the AAL atlas consisting of 90 nodes based on cortical gyri and subcortical nuclei (Tzourio-Mazoyer N *et al*. 2002), and a functionally-based definition was given by the Power atlas with 264 nodes based on meta-analyses of task and rest data (Power JD *et al*. 2011). Notably, the Power atlas does not include nodes in the bilateral hippocampus, bilateral amygdala and bilateral ventral striatum. Since these regions are of particular interest in cognitive and clinical neuroscience, we additionally included these nodes based on previously published coordinates from meta-analyses (Spreng RN et al. 2009; Liu X et al. 2011; Sabatinelli D et al. 2011), thereby increasing the total number of nodes to 270 (one node per region and hemisphere). This expanded Power atlas has also been used in the previous research (Cao H *et al*. 2014; Braun U *et al*. 2015).

Following the previously published procedures (Cao H *et al*. 2014), the mean time series for each node in both atlases were extracted from the preprocessed images. The extracted time series were then corrected for the mean effects of task conditions (for task data), white matter and cerebrospinal fluid signals, and the 24 head motion parameters (i.e. the 6 rigid-body parameters generated from the realignment step, their first derivatives, and the squares of these 12 parameters, (Satterthwaite TD et al. 2013; Power JD et al. 2014)). The residual time series were then temporally filtered (task data: 0.008 Hz high pass, rest data: 0.008-0.1 Hz band pass) to account for scanner noises.

#### Edge definition and network thresholding

The corrected and filtered time series were subsequently used to build a 90 × 90 (AAL atlas) or 270 × 270 (Power atlas) pairwise correlation matrix for each scan using Pearson correlations. The derived correlation matrices were further thresholded into 41 densities ranging from 0.10 to 0.50 with an increment interval of 0.01. At each density, only the connections with correlation coefficients higher than the given threshold were kept as true internode connections in the matrices. The density range was based on common practice in the literature and on empirical data where small-world networks are present within the range (Achard S and E Bullmore 2007; Cao H *et al*. 2014; Cao H *et al*. 2016). Afterwards, edges in binary networks were defined by assigning a value of 1 to the connections that survived a given threshold, and edges in weighted networks were given as the original correlation coefficients of the survived connections. For both binary and weighted networks, a value of 0 was assigned to the connections that did not survive a given threshold. As a result, four adjacency matrices were generated for each scan: 90 × 90 binary matrix, 90 × 90 weighted matrix, 270 × 270 binary matrix and 270 × 270 weighted matrix. Graph theory based brain network measures were subsequently calculated from these derived matrices.

#### Graph theoretical measures for functional connectomes

We computed a series of graph-based connectomic measures that are commonly reported in the literature evaluating the network connectivity strength, network segregation and integration, small-world and modular structures, assortative and hierarchical organizations and nodal centrality. These measures can be generally divided into two categories: global measures and local measures. The global measures quantify the characteristics of brain system as an entity, which include:

1. Mean connectivity: mean of all elements in the correlation matrix;
2. Small-worldness: an index assessing the combination of network segregation (clustering) and network integration (path length);
3. Transitivity: normalized global metric of network clustering;
4. Characteristic path length: average shortest path length between all pairs of nodes in the network;
5. Global efficiency: average inverse of shortest path length between all pairs of nodes in the network;
6. Modularity: degree to which the network can be divided into non-overlapping communities;
7. Number of modules: number of communities the network can be divided into;
8. Assortativity: tendency for nodes to be connected with other nodes of the same or similar degree;
9. Hierarchy: power law relationship between degree and clustering coefficients for all nodes in the network.

Accordingly, the local measures quantify the properties of each network node, including:

1. Node strength: mean connectivity of a given node;
2. Node diversity: variance of connectivity of a given node;
3. Node degree: number of links connected to a given node;
4. Clustering coefficient: proportion of node's neighbors that are also neighbors of each other;
5. Node path length: average path length between a given node and all other nodes in the network;
6. Local efficiency: inverse of shortest path length for a given node;
7. Betweenness centrality: fraction of shortest paths in the network that pass through a given node;
8. Within-module degree: local degree of a given node in its own module relative to other nodes;
9. Participation coefficient: ability of a given node in connecting different modules relative to connecting its own module.

All measures were computed using the Brain Connectivity Toolbox (BCT, https://sites.google.com/site/bctnet/). For a more detailed description of these graph theoretical measures, please refer to the previous publications (Bullmore E and O Sporns 2009; Rubinov M and O Sporns 2010; Bullmore ET and DS Bassett 2011). Of note, the computations for small-worldness and modular partitions were based on 100 network randomizations, and Louvain greedy algorithm was used for the optimization of modularity quality function Q (with resolution parameter ? = 1)(Newman ME 2006; Blondel VD et al. 2008). After computation, all derived connectomic measures were averaged across all densities to ensure that results were not biased by a single threshold.

### Assessment of reliability using generalizability theory

#### Generalizability theory

Details of generalizability theory have been described in the supplementary materials. In brief, generalizability theory is an extension of classical test theory with intra-class correlation coefficients (ICC) as index of reliability (Shrout PE and JL Fleiss 1979; Barch DM and DH Mathalon 2011; Cao H *et al*. 2014). It pinpoints the source of different systematic and random variances by decomposing the total variance into different facets of measurement (Shavelson RJ and NM Webb 1991; Barch DM and DH Mathalon 2011). Here, the total variance of the outcome measures (*σ*^2^(*Xpsd*)) were decomposed into 1) the participant-related variance *σ*^2^(*p*), 2) the scan site-related variance *σ*^2^(*s*), 3) the session-related variance *σ*^2^(*d*), 4) their two-way interactions *σ*^2^(*ps*), *σ*^2^(*pd*), *σ*^2^(*sd*), and 5) their three-way interaction and random error *σ*^2^(psd,e)(Shavelson RJ and NM Webb 1991; Noble S et al. 2016).

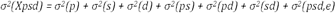

The reliability coefficients are then calculated in terms of the participant-related variances and variances of interest, which are analogous to the ICC values derived from classical test theory (Shavelson RJ and NM Webb 1991). Similar to ICC estimates, the reliability coefficients in generalizability theory include G-coefficient 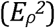 and D-coefficient 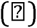, which evaluate relative consistency and absolute agreement of the target measures, respectively (Shavelson RJ and NM Webb 1991).

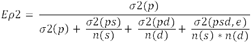

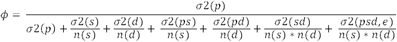

where *n(i)* represents the number of levels in factor *i*. According to established criteria (Shrout PE and JL Fleiss 1979; Cao H *et al*. 2014; Forsyth JK et al. 2014), both G- and D-coefficients can be interpreted as: poor reliability (< 0.4); fair reliability (0.4 − 0.59); good reliability (0.6 – 0.74); and excellent reliability (> 0.74).

Generalizability theory can be applied to two types of studies, namely, the generalizability study (G-study) and the decision study (D-study). In the G-study, the reliability coefficients are estimated based on the facets and their levels in the studied sample (here n(s) = 8, n(d) =2)(Shavelson RJ and NM Webb 1991; Forsyth JK et al. 2014), while in the D-study, the researchers define the universe they would like to generalize into, which may contain some or all of the facets and levels in the overall universe of observations (Shavelson RJ and NM Webb 1991; Noble S et al. 2016). Since in a neuroimaging “big data” context, a “nested” design is commonly used where each participant is scanned once only at one site and different subjects could be scanned on any number of different scanners, the expected site- and session-related variances would be higher than those in a balanced design as used in this study (Lakes KD and WT Hoyt 2009). We therefore recomputed the reliability coefficients to generalize our results to n(s) = 1 and n(d) = 1, which correspond to the expected reliability in a “nested” design with distinct subjects between sites and sessions.

#### Statistics

We performed both G- and D-studies on each of the examined graph properties. For each study, the measurement variances were decomposed using a three-way analysis of variance (ANOVA) model, where graph properties were entered as dependent variables and subject, site and session were entered as random-effect factors. The estimated variances for each factor were then subjected to the reliability coefficients formula, and G- and D-coefficients for each of the graph properties were calculated. This procedure was repeated for each processing scheme and each paradigm.

We then used the resulting coefficients to explore several scientifically interesting questions. In particular, we asked 1) whether there were significant reliability differences between global and local properties; 2) whether different processing schemes (i.e. AAL binary, AAL weighted, Power binary, Power weighted) resulted in significant differences in reliability measures; and 3) whether different fMRI paradigms (i.e. WM, EM encoding, EM retrieval, EP, RS) generated similar reliability. Here, a repeated-measures ANOVA was employed to answer these questions, where reliability measures for each property were given as dependent variable, processing scheme and fMRI paradigm were set as within-subject factors, and property type (global and local) was set as a between-subject factor. The main effects for each of the factors were then estimated.

#### Node-wise reliability in D study

Given the fact that the D study yielded significantly lower mean reliability than the G study (see Results), particularly for measures of nodal centrality (i.e. degree, betweenness, within-module degree, participation coefficient), we further probed the reliability of centrality measures for each node in the D study, in order to ascertain the most reliable nodes in the brain in different paradigms. Here, we only utilized weighted networks to maximize reliability and minimize confounds, since binary networks were shown to be significantly less reliable than weighted networks (see results). The reliability computation followed the same procedure as described above.

### Comparison of statistical power between multisite and single-site studies

The ultimate goal of performing a multisite study is to boost statistical power. However, since multisite studies introduce variance related to different scanners, which is not the case with single-site studies, the resulting decrease in reliability would likely lead to loss of power. For this reason, we further estimated the minimal sample size that is required for a multisite study to achieve comparable power to that of a single-site study. Here, we first probed reliability differences of connectomic measures between a multi-site study and a single-site study where all subjects were scanned on the same scanner. Using D study formula, the G coefficients of all examined measures were recalculated for each of the eight sites. For each site (16 scans with 8 subjects and 2 sessions), three variance components were considered: subject, session and subject 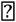 session. The reliability coefficients were computed in terms of these three components and then averaged across all eight sites. By this procedure we acquired the empirical estimates evaluating the reliability for a single-site study.

In a situation of perfect reliability (r = 1), the effect size of measurement equals its “true” effect size. However, the effect size attenuates when the reliability of measurement decreases. Therefore, low reliability would bias the “true” effect size of the measurement and in turn lead to power loss (Cohen J 1988). Here, we calculated the empirical effect sizes for all graph measures based on their reliability coefficients in multisite and single-site study context. We considered a set of “true” effect sizes ranging from 0.3 to 0.9 (interval of 0.2) to mimic different levels of effect size in a case-control study (small: 0.3; medium: 0.5; large: 0.7; very large: 0.9; (Cohen J 1988)). The effect sizes for each measure were computed according to Cohenʼs formula (Cohen J 1988):

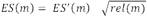

where *ESʼ(m)* is the “true” effect size of the given measurement, *rel(m)* is the reliability of the measurement, and ES(m) is the derived effect size under that reliability estimate. The percent changes of effect size between multisite and single-site studies were subsequently acquired for each measurement (see Table S15).

The power estimations were performed using the R statistical power analysis toolbox pwr (https://cran.r-project.org/web/packages/pwr/index.html). Here, for each of the “true” effect sizes, we calculated statistical power across a range of sample sizes for both multisite and single-site studies. This generated a set of functions depicting the relationship between statistical power and sample size for both studies and thus provided the information on the optimal sample sizes for each measure in a multisite study context.

### Comparison of different data aggregation approaches

We further addressed another practical question: at which level should we aggregate data from separate runs of a paradigm? For example, one could merge the outcomes by computing the connectomic measures for each scan run separately and then averaging the derived measures from multiple runs. Alternatively, one could merge the original data by concatenating time series from multiple runs and then computing the connectomic measures from the concatenated time series. We refer the former as the “merging results” approach and the latter as the “merging raw data” approach. Previous research has shown that concatenation of time series using single-session data dramatically decreases reliability (Cao H *et al*. 2014), suggesting that “merging raw data” may not be an optimal choice for data aggregation. However, by using single-session data, splitting and concatenation of time series would also lead to the decrease of number of time points, bringing difficulty in the interpretation of the observed reliability changes. Therefore, it would be important to investigate whether the same reliability results apply to data concatenation by using scans from multiple sites and/or sessions. Since concatenation of multiple scans would dramatically increase the number of time points and thus benefit reliability, any reliability reductions in the context of merged data would be most likely due to the concatenation method itself.

Here, we aimed to give a direct comparison of reliability measures derived from “merging results” and “merging raw data” approaches, in order to inform a superior data aggregation approach in a multisite, multisession study. The EP task used in this study offered an opportunity to explicitly investigate this question since it comprised two consecutive identical runs on each scan day (5.5 min each, see text above). Here, by “merging raw data” the preprocessed time series of both runs were concatenated and brain graph measures were computed from the concatenated time series (i.e. 11 min). In contrast, by “merging results” the graph measures were computed for each run separately and then averaged to acquire the mean measures for both runs. We subsequently calculated the reliability coefficients for the resulting measures from both approaches. A repeated-measures ANOVA model was employed to compare the reliability differences between these two approaches and single session, with the processing approach as within-subject factor and reliability measures as dependent variable.

## Results

### Brain graph reliability for each paradigm

Overall, in the context of the 8-site, 2-session study, we observed fair to excellent reliability for almost all computed measures in all paradigms, regardless of processing scheme (Fig. 2A-2E, Fig. S1A-S1E). The only exceptions for this were measures of assortativity and hierarchy during resting state, two second-order metrics that showed relatively poor reliability using the AAL atlas 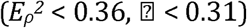 and fair reliability using the Power atlas 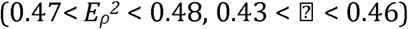 Among the remainder, the most reliable measures were mean connectivity 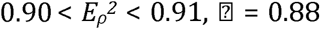 transitivity 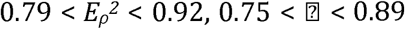 global efficiency 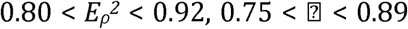 and node strength 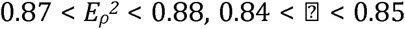 In addition, small-worldness, local efficiency, path length, clustering coefficient and modularity also showed excellent reliability when using the Power atlas and weighted networks 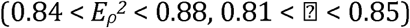 (Table S3).

**Fig. 2.**
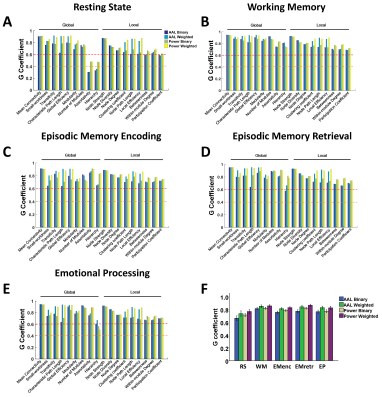
The reliability outcomes (G coefficients) for examined graph properties, paradigms and processing schemes in the G study context, wherein N_sites_=8 and N_sessions_=2 (panel A: resting state; panel B: working memory; panel C: episodic memory encoding; panel D: episodic memory retrieval; panel E: emotional processing). Panel F shows the statistical comparison between different paradigms and schemes by averaging all graph properties. The orange dashed lines indicate the level of fair reliability 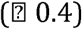 and the red dashed lines indicate the level of good reliability 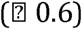.

The reliability of brain graphs constructed from scans involving cognitive tasks showed a general advantage compared with those from the resting state (Fig. 2B-2E, Fig. S1B-S1E). In particular, all computed measures with all processing schemes demonstrated good to excellent reliability in terms of G coefficients for the working memory and episodic memory encoding paradigms 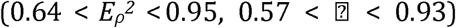Similar results also applied to the episodic memory retrieval and emotional processing paradigms 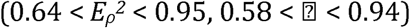with hierarchy as the only exception with fair to excellent reliability 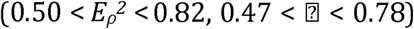Notably, when using Power weighted networks, almost all measures from all tasks showed excellent reliability 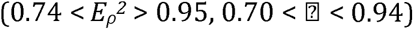 The only exceptions here were hierarchy and modular numbers, with fair to good reliability during working memory and emotional processing 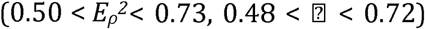and good reliability during episodic memory encoding 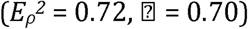respectively (Tables S5, S7, S9, S11).

In the D-study with N_sites_=1 and N_sessions_=1, we found dramatically reduced reliability compared with G-study, particularly when analyzed with binary networks 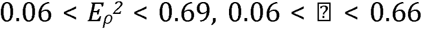for all paradigms, Fig. 3A-3E, Fig. S2A-S2E). Nevertheless, a few measures still showed fair to good reliability across all paradigms and across both atlases, though only for weighted networks. These included measures assessing network connectivity strength 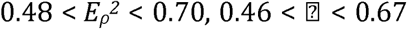and network segregation and integration 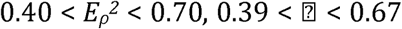 (Tables S4, S6, S8, S10, S12). Other measures, such as small-worldness and number of modules, also reached fair to good reliability in the working memory, episodic memory retrieval and emotional processing tasks 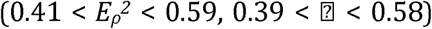Together, these findings suggest that measures of network connectivity and network segregation and integration are the most reliable measures in human functional connectomics, even when pooling data in which different subjects are scanned once on different scanners.

**Fig. 3.**
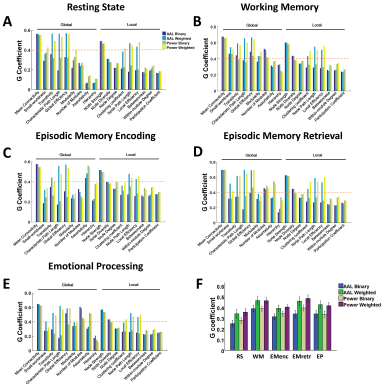
The reliability outcomes (G coefficients) for examined graph properties, paradigms and processing schemes in the D study context, wherein N_sites_=1 and N_sessions_=1, simulating the design of the typical “big data” application involving pooling of data from different scanning sites (panel A: resting state; panel B: working memory; panel C: episodic memory encoding; panel D: episodic memory retrieval; panel E: emotional processing). Panel F shows the statistical comparison between different paradigms and schemes by averaging all graph properties. The orange dashed lines indicate the level of fair reliability 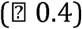

### Reliability comparisons between paradigms, processing schemes and property types

For both G- and D-studies, the results revealed that fMRI paradigms (F > 11.00, P < 0.001) and processing schemes (F > 8.14, P < 0.001) significantly influenced reliability coefficients (Figs. 2F, 3F). Specifically, graph measures computed from all cognitive tasks showed higher reliability than those from resting state (F > 5.05, P < 0.04). Within cognitive tasks, working memory and episodic memory retrieval paradigms showed higher reliability than episodic memory encoding and emotional processing paradigms (F > 9.49, P < 0.007). In terms of processing schemes, both AAL weighted and Power weighted networks had higher reliability than AAL binary and Power binary networks (F > 8.22, P < 0.01), suggesting that the use of weighted networks would increase reliability in multi-center functional connectomics studies. In contrast, property types (global and local) did not show significant effects on reliability coefficients (F < 1.44, P > 0.25), suggesting that global and local properties are equally reliable.

### Variance components of functional graph measures

We report the results of variance isolation derived from the Power weighted networks here, since this scheme in general yielded the highest reliability (Figs. 2-3). Overall, for all properties in all paradigms, the three largest variance components were participant, participant 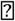 site and participant 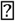 site 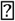 session (Figs. S3-S7). These three components together accounted for more than 80% of the total variance for almost all properties. In contrast, session-related variance was the smallest, less than 1% of the total variance for most properties. In addition, site-related variance was much smaller (2 to 15 times) than participant-related variance for all properties. These results indicate that between-subject variance is much larger than within-subject variance in functional connectomics studies using multisite, multisession data, a property that makes graph theoretical analysis useful for big data applications.

For resting state, the proportion of variance attributed to participant ranged between 9% and 49%, with the highest proportion in relation to global efficiency and lowest in relation to hierarchy. For the cognitive tasks, the participant variance ranged between 21% and 64%. Here, the property with the highest participant-related variance was mean connectivity, which accounted for about 60% of the total variance for each task. Other properties with high participant-related variance included measures of network segregation and integration (path length, transitivity, clustering coefficient, efficiency) and node strength, which in general accounted for around 50% of the total variance for each task. In contrast, participant-related variance represented a low proportion (less than 40% of the total) for measures of small-worldness, modularity, hierarchy and nodal centrality (degree, betweenness, within-module degree, participation coefficient). These results suggest that graph properties evaluating network segregation and integration are more trait-related measures, while those evaluating small-world organization, modular structure and centrality are more state-related measures.

### Node-wise reliability in D study

We found that cognitive tasks had considerably more reliable nodes than resting state for all centrality measures with both AAL and Power atlases (Fig. 4). Interestingly, the reliable nodes highly overlapped between the four cognitive paradigms. With both AAL and Power atlases, the reliable nodes 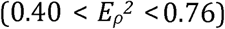mainly mapped to the fronto-parietal system (e.g., superior, middle and inferior frontal gyri, superior and inferior parietal lobules), default-mode system (e.g., medial frontal cortex, angular gyrus, precuneous, superior and middle temporal gyri, temporal pole), visual system (e.g., superior and middle occipital gyri, fusiform gyrus, calcarine sulcus, cuneous), limbic system (e.g. cingulate cortex, orbitofrontal cortex, parahippocampal gyrus, amygdala, hippocampus, insula), sensori-motor system (e.g., precentral gyrus, postcentral gyrus, supplementary motor area, paracentral lobule) and subcortex (e.g., caudate, pallidum, thalamus)(Fig. 4, Tables S13,S14).

**Fig. 4.**
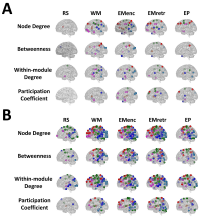
The most reliable nodes in terms of nodal centrality measures in the D study context, wherein N_sites_=1 and N_sessions_=1, simulating the design of the typical “big data” application involving pooling of data from different scanning sites (panel A: AAL weighted networks; panel B: Power weighted networks). Note that the default-mode (blue), visual (cyan) and sensori-motor (green) systems showed high reliability in both resting state and cognitive tasks, while the fronto-parietal (red), limbic (magenta) and subcortical (orange) systems were predominantly reliable in cognitive tasks (see Table S13, S14 for details). Abbreviations: RS = resting state; WM = working memory; EMenc = episodic memory encoding; EMretr = episodic memory retrieval; EP = emotional processing.

In contrast to the results for task paradigms, the resting state in general showed fewer reliable nodes, in particular when using the AAL atlas. Here, hardly any nodes had a G coefficient 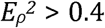for all centrality measures (Fig. 4, Table S13), indicating poor feasibility of aggregating “nested” data for the investigation of nodal centrality in resting state with the AAL atlas. In terms of the Power atlas, the reliable nodes 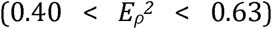 were mainly distributed in the default-mode system (e.g., medial frontal cortex, angular gyrus, precuneous, superior and middle temporal gyri), visual system (e.g., middle and inferior occipital gyri, lingual gyrus), and sensori-motor system (e.g., precentral gyrus, postcentral gyrus, supplementary motor area, paracentral lobule) (Table S14). Notably, these systems were part of the reliable systems found in cognitive tasks, suggesting that the reliability distribution of nodal centrality consists of a set of systems that is independent of active tasks and another set of systems that is reliable only when tasks are presented.

### Comparison of statistical power in multisite versus single-site studies

The reliabilities of all connectomic measures were substantially higher in the single-site study compared with the multisite study context. As in the multisite study, measures of connectivity strength, network segregation and integration had highest reliability of all measures in the single-site study. In addition, measures of hierarchy, modular structure, node diversity and centrality were considerably more reliable in the single-site study than in the multisite study (Table S15).

The power analysis revealed that with a small effect size (d = 0.3), a considerably large total sample size (600 to 1000) was required for all examined properties to gain an adequate level of power (i.e., ≥ 0.8) (Figs. 5, S8). However, with an increase of effect size, the sample size required to achieve adequate power was dramatically decreased. For the most reliable measures including network connectivity, network segregation and integration, only 200 to 250 subjects in total were needed to detect a medium effect (d = 0.5), only 100 to 150 to detect a large effect (d = 0.7), and only < 100 to detect a very large effect (d = 0.9) (Fig. 5).In contrast, for relatively less reliable measures such as hierarchy and nodal centrality, a total sample size of > 150 was required even with a very large effect size (d = 0.9).

**Fig. 5.**
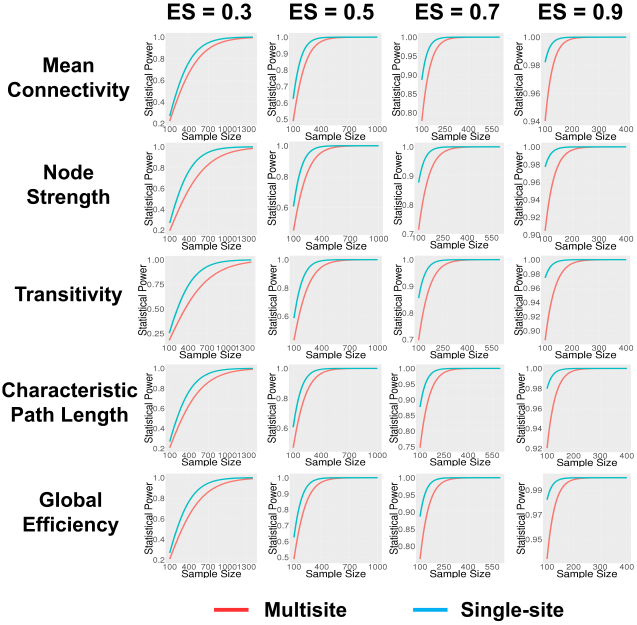
Statistical power as a function of total sample size across multiple effect sizes for five selected connectomic measures. The red lines represent power for multisite studies while the blue lines represent power for single-site studies, based on Cohen’s d for two-tailed contrast of two independent groups at 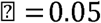= 0.05. The effect sizes have been adjusted downward for observed reliabilities of each connectomic measure in the multisite and single-site contexts, respectively. Although higher levels of power are achieved with smaller sample sizes in the single-site compared with multisite context, multisite studies achieve acceptable levels of power (≥ 0.8) with moderate to large effect sizes (ES ≥ 0.5) beginning at sample sizes of approximately 200 subjects. See Fig. S8 for comparable analyses for the remaining connectomic measures.

### Comparison of data pooling methods

By merging results from both runs, reliability of all studied graph measures increased compared with those derived from a single scan (Figs. S9-S10). In contrast, the “merging raw data” approach reduced reliability for almost all measures. A direct comparison between the two methods demonstrated a significant difference in graph reliability, where “merging results” approach yielded a significantly higher reliability than a single run (t > 4.31, P < 0.001) and “merging raw data” approach (t > 3.93, P < 0.001) in both G- and D-studies. Moreover, graph measures derived from single run showed significantly higher reliability than those derived from “merging raw data” approach in both G- and D-studies (t > 2.28, P < 0.04). These results support the superiority of using a “merging results” approach in data aggregation in the study of human functional connectomes.

## Discussion

This study investigated a fundamental question in human functional connectomics research: how feasible is it to aggregate multisite fMRI data in large consortia? Our results demonstrated that 1) the connectomic measures derived from different sites and sessions showed generally fair to good reliability, particularly for measures of connectivity strength, network segregation and network integration; 2) the reliability of connectomic measures varied between different fMRI paradigms, with cognitive tasks significantly more reliable than resting state; 3) choice of processing schemes significantly influenced reliability, with weighted networks more reliable than binary networks; 4) the most reliable nodes in the brain differed between rest and tasks; 5) a total sample size of 250 participants or more was sufficient for case-control studies in multisite functional connectomics research if the effect size of group differences is at least moderate (≥ 0.5); and 6) different data aggregation approaches significantly affected outcomes, with “merging results” approach superior than “merging raw data” approach. These results provide direct evidence for the feasibility of pooling large sets of functional data in human connectomics and provide useful guidelines for sample size, data analysis approaches, and aggregation methods that equate with adequate levels of reliability and statistical power.

### Overall reliability of functional graph measures

The graph modelling of human connectomes quantifies a series of topological measures evaluating the organization of the brain system. This typically includes measures for network connectivity, network integration and segregation, small-world and modular structures, assortative and hierarchical organizations, and node centrality (see (Bullmore E and O Sporns 2009; Rubinov M and O Sporns 2010) and Methods). Here, using our multisite, multisession data, we found that the vast majority of these measures were reasonably reliable during both resting state and cognitive tasks. This finding is in line with previous studies using test-retest data, where fair to excellent test-retest reliability have been shown for functional graph metrics in various fMRI paradigms, including resting state (Braun U *et al*. 2012; Cao H *et al*. 2014; Welton T et al. 2015; Termenon M *et al*. 2016), working memory (Cao H *et al*. 2014), emotional processing (Cao H *et al*. 2014) and attentional control (Telesford QK et al. 2010). These prior publications and our present data suggest that the graph theory based connectomic measures are associated with overall high between-subject variance and relatively lower within-subject variance. Indeed, our analyses revealed that subject-related variance was one of the largest components for almost all examined properties in all paradigms. In contrast, within-subject variance such as scan site- and session-related variance, were found to be much smaller than subject-related variance, suggesting the feasibility of using graph based measures in the study of human functional connectomes in large consortia. Notably, two other relatively large components were the subject 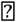 site and subject 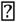 site 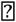 session interactions. This pattern is also consistent with previous findings in brain activity measures during working memory (Forsyth JK et al. 2014) and emotional processing (Gee DG et al. 2015). These results suggest that many fMRI measures are sensitive to factors such as subject alertness, diurnal variations, situational distractions, and variations in head placement, among others (Meyer C et al. 2016). Because these factors were not controlled in our traveling subjects study, and indeed are difficult to homogenize across sites and time, the reliability coefficients presented here likely represent a lower bound estimate of reliability (i.e., theoretically higher reliabilities would be obtained with greater standardization of time of day, head positioning, subject alertness, etc.).

Interestingly, some of the examined graph properties were associated with particularly high proportions of subject-related variance during the cognitive tasks, including measures of connectivity strength (e.g., mean connectivity, node strength), network segregation (e.g., clustering coefficient, transitivity) and network integration (e.g., path length, global/local efficiency). For these measures, approximately 50%-60% of total variances were attributed to subjects, indicating that subject-related variance was larger than any other components. As a result, higher reliabilities have been detected for these metrics compared with others. When generalized to the context in which different subjects are scanned once on different scanners, fair to good reliabilities were still evident for these properties. Since a similar pattern has also been reported in the previous test-retest studies (Telesford QK et al. 2010; Cao H *et al*. 2014; Termenon M *et al*. 2016), these results suggest that graph properties for connectivity strength and network segregation and integration are particularly robust across scan sites and scan sessions and are thus possibly reflective of stable, participant-specific features of brain organization. In contrast, measures assessing the small-world and modular structures (e.g., small-worldness, modularity, number of modules) and nodal centrality (e.g., degree, betweenness, within-module degree, participartion coefficient) had in general equal proportions of subject-related and error-related components, suggesting that these measures are to a greater degree sensitive to within-subject factors and thus potentially more state-related. Prior work has shown that measures of network segregation and integration are highly heritable (Smit DJ et al. 2008; Fornito A et al. 2011), while measures of small-world and modular structures are dynamic during different behavioral and cognitive states such as finger tapping (Bassett DS et al. 2006), motor learning (Bassett DS et al. 2011; Bassett DS et al. 2015) and memory (Braun U *et al*. 2015). These findings converge with our results in suggesting that trait- and state-related characteristics of brain functional systems may be captured by different graph based connectomic measures.

### Effects of fMRI paradigms on reliability

We observed a significant effect of fMRI paradigm on graph reliability. Specifically, cognitive tasks were observed to produce more reliable network diagnostics than resting state in our study. This is in agreement with a previous finding that graph properties derived from a working memory task exhibited higher test-retest reliability than those from resting state (Deuker L l*et al*. 2009). Notably, while the vast majority of reliability studies to date have focused on resting state, little research has been done on the direct comparison of graph reliability between rest and tasks (Welton T et al. 2015). The only two studies available in the literature, however, have drawn different conclusions (Deuker L l*et al*. 2009; Cao H *et al*. 2014). While it is still an open question whether resting state or cognitive tasks would yield higher reliability, our present data have provided new evidence that connectomic measures computed from tasks are more reliable than those from rest. This pattern may relate to at least two factors. First, compared to the “mind-unconstrained” resting state, subjects' behaviors and cognitions are well controlled by the given task paradigms, which require considerably more attentional effort. Second, since resting state involves the engagement of internally focused thoughts such as introspection, future envisioning and autobiographical memory retrieval (Buckner RL et al. 2008), it is by nature more vulnerable to state-related within-subject factors such as mood, tiredness, diurnal variations, and scan environment. In addition, another possible interpretation for the higher reliability of cognitive tasks would be the generally longer time series in tasks compared with rest. To keep the integrity of task time series, we did not arbitrarily discard any task-related time points to match the length of resting state, which may to some degree confound our results. However, given the evidence that the detected reliability differences between rest and tasks were qualitatively the same in length-matched and length-unmatched time series in previous studies (Deuker L l*et al*. 2009; Cao H *et al*. 2014), it is highly unlikely that our results are mainly driven by this discrepancy.

Another interesting finding between paradigms is the significant reliability differences between cognitive tasks. Here, the working memory and episodic memory retrieval tasks are significantly more reliable than episodic memory encoding and emotional processing tasks. This finding is not too surprising given that working memory paradigms have repeatedly been shown to be more reliable than emotional processing (Plichta MM *et al*. 2012; Cao H *et al*. 2014; Forsyth JK et al. 2014; Gee DG et al. 2015), possibly due to strong habituation effects in the emotional tasks (Breiter HC et al. 1996; Johnstone T et al. 2005; Plichta MM *et al*. 2012) and additional attentional efforts in the more demanding memory tasks (Cao H *et al*. 2014). The higher reliability of episodic memory retrieval compared with encoding has also been previously reported in terms of fMRI activation measures (Clement F and S Belleville 2009). This reliability difference may relate to the choice of performance strategies and intrinsic learning effects associated with the encoding paradigm, which renders the encoding phase less reliable than the retrieval phase.

### Effects of processing schemes on reliability

Our study also found a significant effect of processing scheme on the reliability of connectomic measures. Specifically, weighted networks were more reliable than binary networks using both AAL and Power atlases, particularly in the D study context. Unlike binary networks that have simplified network structures by setting the strength of all edges with the same value of one, weighted networks preserve the original information on network connectivity strength. As a result, weighted networks can characterize the real network topology more precisely and thus promote reliability. This finding encourages the use of weighted networks instead of binary networks in functional connectomic studies with big data. In addition, although previous research has identified a significant difference in graph reliability between AAL and Power atlases (Cao H *et al*. 2014), we failed to replicate this finding. Notably, although a trend towards higher reliability with the Power atlas was seen in binary networks (Figs 2F, 3F) which is consistent with the prior finding, this difference was not observed in the weighted networks. Together, our results suggest that both anatomic and functional atlases are feasible in functional connectomic research when weighted networks are employed.

### Reliable nodes in rest and tasks

Across both resting state and cognitive tasks, the reliable nodes were predominantly distributed in the default-mode, visual and sensori-motor systems. This result is highly parallel to the results of a recent study using the Human Connectome Project (HCP) test-retest data (Termenon M *et al*. 2016), where exactly the same distribution was reported for the resting state. Interestingly, all these three systems serve as the primary functional systems in the human brain. Default-mode network is involved in brain's resting state and becomes active when individuals are focused on internal thoughts (Buckner RL et al. 2008). The visual system is directly associated with the visual functioning of the experimental paradigms, and the sensori-motor system may relate to the motor response during active tasks and the sensation of environment changes during resting state. The function of these systems makes them plausible to be more robust than other systems independent of fMRI paradigms.

Besides the above systems, the cognitive tasks also showed high reliability for nodes in the fronto-parietal, limbic and subcortical systems. Notably, these systems are pivotal to human cognitive functions such as memory (Prabhakaran V et al. 2000; McNab F and T Klingberg 2008) and emotion recognition (Phillips ML et al. 2003) and are strongly associated with the memory-emotion task battery used in this study (Forsyth JK et al. 2014; Gee DG et al. 2015). Together, these findings suggest that the cognitive tasks would increase the reliability of the multimodal cognitive-emotional systems, while the primary functional systems are consistently robust through different brain states/imaging paradigms.

### Statistical power for multisite and single-site studies

By calculating reliability for each single site, we found that the reliability distribution followed the same pattern as that in the multisite study, where measures of connectivity strength, network segregation and integration showed highest reliability. This finding is in good agreement with previous studies using single-site data (Telesford QK et al. 2010; Cao H *et al*. 2014; Termenon M *et al*. 2016). Notably, there was the least amount of drop-off in terms of comparative reliability of these measures when comparing the single site study to the multisite study context. In contrast, measures with relatively low reliabilities became considerably less reliable when using the multisite design, suggesting that these lower reliability measures are more vulnerable to loss of power in a multisite study.

The follow-up power analysis demonstrated that, although a considerably larger sample size is required to compensate for power loss in a multisite study when the effect size is small, with medium to large effects, sample sizes required for adequate to excellent power are in the range typical of consortium studies (i.e., total N of 200 to 500). In particular, for measures with relatively high multisite reliability (i.e., connectivity strength, network segregation and integration), the sample size needed for 80% power can be as small as 250 for a medium effect, 150 for a large effect, and approximately 50 for a very large effect. Interestingly, a recently study using simulated data also found that approximately 120 subjects per group are sufficient to yield satisfying power for network connectivity measures in multisite studies (Dansereau C et al. 2017). Since a total sample size larger than 250 subjects is increasingly common in large consortia, we conclude that most functional connectomic studies using big data are likely to be reasonably well-powered.

### Effects of data pooling methods on reliability

By comparing two data pooling approaches, we found that “merging results” is associated with significantly better reliability of brain graph measures compared with the “merging raw data” approach. This result is consistent with previous work using single-session data that also revealed a significant decrease of graph reliability by the chopping and concatenation of fMRI time series (Cao H *et al*. 2014). Notably, our current finding was derived from the concatenation of data from two identical runs, which increased the total number of time points by a factor of two. Considering this, the reliability change reported here is most likely induced by the “concatenation” approach itself rather than the loss of data points. While speculative, the poor performance of “concatenation” approach may relate to the modification of the fundamental characteristics of original fMRI series such as signal frequency, which renders the concatenated signals particularly sensitive to physiological noise and other artifacts (Gavrilescu M et al. 2008). In contrast, the average of graph properties computed from separate runs significantly increased reliability. This result is intuitive since the mean calculation of multi-run data statistically reduces run-specific noise and thus boosts reliability.

### Limitations

We acknowledge several methodological limitations for our study. First, although we have constructed brain networks with several different processing schemes in this work, the reliability estimates computed in this study are still dependent upon the methodological choices that were not varied in the processing stream, such as the preprocessing pipeline (Braun U *et al*. 2012), filter frequency (Deuker L l*et al*. 2009; Braun U *et al*. 2012) and selected thresholds (Schwarz AJ and J McGonigle 2011; Termenon M *et al*. 2016). Second, while we have provided data on a set of commonly used fMRI experiments evaluating the cognitive, emotional and resting functions of the brain, our results are nevertheless influenced by the employed paradigms, and it is possible that results could vary with other task paradigms. Third, our reliability study is based on a balanced design where each subject is evaluated at all of the different sites across two sessions. This is different from a more commonly used “nested” design in which different subjects are evaluated on different scanners. We sought to generalize our results to mimic this situation using the D-study extension, where we are essentially modeling how well one scan randomly sampled from among the set of 16 scans available for each subject reflects their “true” score for each graph property. Fourth, except for several basic factors such as demographics, head motion and task response rate, we did not deliberately perform a strict quality control on our data. This is to mimic the situation in large consortia in which data from different sites and sessions may not be well balanced and thus can be influenced by a set of physiological, psychological and neuroimaging factors. Since these confounds would generally increase variances unrelated to subject and thus decrease the outcome reliability (Gorgolewski KJ et al. 2013), our results are likely to underestimate the reliability that would obtain in a perfectly matched dataset. Fifth, given that sample size has a significant effect on reliability estimates (Termenon M *et al*. 2016), the functional connectomic measures would potentially be even more reliable in larger datasets such as the HCP (Van Essen DC *et al*. 2013) and the 1000 Connectome Project (Biswal BB *et al*. 2010) compared to our sample of 8 traveling subjects. Last but not least, while our results are derived from a group of healthy participants, further investigations are still warranted to test these reliability findings in clinical populations.

## Conclusions

In conclusion, this multisite, multisession reliability study has revealed overall fair to good cross-site reliability for a majority of graph based functional connectomic measures during both resting state and cognitive tasks, with particularly high reliability for measures of connectivity strength, network segregation and network integration. The detected reliabilities are influenced by fMRI paradigms and network construction schemes, with higher reliabilities detected for graphs generated from task-based fMRI and weighted networks. Moreover, this study identified a differential distribution of most reliable brain nodes in resting and task-based data, and provided empirical evidence that satisfactory statistical power can be acquired with a total of 250 subjects in a multisite study when the effect size is medium. Finally, we demonstrated the superiority of a “merging results” approach compared with a “merging raw data” approach in terms of data aggregation. Our findings offer direct evidence for the feasibility of pooling large set of fMRI data for both rest and tasks in large consortia and encourage the use of multisite, multisession scans to promote the power for human functional connectomics.

## Acknowledgements

This work was supported by the National Institute of Health grants U01 MH081902 to Dr. Cannon, P50 MH066286 to Dr. Bearden, U01 MH081857 to Dr. Cornblatt, U01 MH82022 to Dr. Woods, U01 MH066134 to Dr. Addington, U01 MH081944 to Dr. Cadenhead, R01 U01 MH066069 to Dr. Perkins, R01 MH076989 to Dr. Mathalon, U01 MH081928 to Dr. Seidman, and U01 MH081988 to Dr. Walker.

